# Significant and persistent carryover effects in Scots pine

**DOI:** 10.1101/2024.08.23.609343

**Authors:** Annika Perry, Joan K. Beaton, Jenni A. Stockan, Glenn R. Iason, Joan E. Cottrell, Stephen Cavers

**Affiliations:** UK Centre for Ecology & Hydrology (UKCEH), Bush Estate, Midlothian, EH26 0QB, UK; Ecological Sciences, James Hutton Institute, Craigiebuckler, Aberdeen, AB15 8QH, UK; Northern Research Station, Forest Research, Roslin, EH25 9SY, UK

## Abstract

After tracking Scots pine plants from germination to 15 years old, through contrasting early life environments, we observed significant and persistent carryover effects. Groups of plants from common genetic backgrounds were raised in distinct nursery environments, and growth and phenology traits were measured repeatedly once trees had established in their field sites. Growth and phenology differences were evident for 10 and 6 years post-transplantation to the field, respectively. There was a clear interaction between site of origin and carryover effect, indicating that local adaptation also played a role. Given the increasing rate of tree planting initiatives being undertaken around the world in the name of the climate and biodiversity crises, and the strong dependence on those initiatives of nursery-grown plants, our finding of strong carryover effects of the early life environment has significant implications.

## Introduction

The environment a plant experiences in its early life can have extremely large effects on its growth and development. However, the extent to which this early influence persists into later life stages, particularly following transplantation from nursery to the field, is less well understood. This phenomenon is sometimes termed a ‘carryover effect’ and has been defined as “*any situation in which an individual’s previous history and experience explains their current performance in a given situation”* (O’Connor et al., 2014). Carryover may be due to movements between environments, such as from a nursery to the field, but equally may occur where the environment at a site changes significantly over the lifetime of a plant. This is likely to have a particular impact on perennial species such as trees (Duputié et al., 2015), which live long enough to experience substantial environmental change throughout their lifetimes. Acclimation, the capacity of an individual to adapt to changes in their environment during their lifetime (Kleine et al., 2021), and the associated long-term shifts in phenotype which result, are components of individual plasticity (Auge et al., 2017; Rohde and Junttila, 2008). In natural settings, especially when establishment is by natural regeneration, plants should develop phenotypes optimised to their environments but may then adjust through acclimation to comparatively gradual changes. In contrast, plants being traded or raised for planting are likely to move abruptly between two or more distinct environments before final transplantation to the field, all within a few years. In the latter context, whilst acclimation may occur, phenotypic development in a prior environment may either impose a cost or provide an advantage in the new environment.

The use of nurseries to raise plants for commercial use is ubiquitous. The protected environment of the nursery allows seedlings to be produced in volume, maintained and treated in a uniform manner (Krasowski, 2009) and, in some circumstances (Osorio et al., 2003), enables direct or indirect selection prior to transplantation. By design, the environments imposed by nurseries directly affect plant growth and development but may also indirectly affect other adaptive traits such as variation in susceptibility to pests and pathogens (Gruffman et al., 2012; Heiskanen and Rikala, 2000; Selander and Immonen, 1992; Villar-Salvador et al., 1999). Several studies have reported carryover effects in plants following abiotic stresses such as salinity (Zandt and Mopper, 2002), drought (Backhaus et al., 2014; Tombesi et al., 2018) and physical disturbance (Gagliano et al., 2014), while numerous studies have focused on plant defence priming in response to biotic challenge (reviewed by Conrath et al., 2002) and more recently by Mauch-Mani et al., 2017). Although the carryover phenomenon has rarely been studied experimentally for trees, examples include: increased water availability affecting transpiration in *Pinus taeda* (Ewers et al., 1999); ozone altering new root growth and carbon allocation in *Pinus ponderosa* (Andersen et al., 1997); nursery environment impacting height and phenology after transplantation in *Betula papyrifera* (Dhar et al., 2015, 2014) and in *Picea abies* and *Larix decidua* (Gömöry et al., 2015).

Plant populations that have been exposed to sufficiently contrasting environments over multiple generations will diverge in fitness-related trait means, reflecting local adaptation to their environment (Leimu and Fischer, 2008). Therefore, despite the obvious benefits conferred on nursery grown seedlings compared to those in the field, it is expected that seedling growth and development could be affected if grown in environments that differ significantly from those at their site of origin (Campbell and Sorensen, 1984). In recognition of the effects of local adaptation, the use of locally sourced seed is an explicit requirement when planting for both conservation and commercial purposes within the natural distribution of a species in the UK (Herbert et al., 1999). However, despite the guidance surrounding the use of local provenances for planting, experimental studies consistently find that the influence of environment on trait variation is almost always the largest effect (Boyer, 1982; Rosique-Esplugas et al., 2021) and the early environment is expected to be particularly highly influential. For example, seedlings from common sources grown in different nursery environments also show differences in their growth (Randall and Johnson, 1998) and phenology (Dhar et al., 2015; Heide, 1993; Westergaard and Eriksen, 1997) after planting.

Carryover effects of the nursery environment might be expected for growth traits due to the impact of early conditions on provisioning (Gömöry et al., 2015) but how long this effect persists is not well-studied. Moreover, long-term experimental testing for this effect is lacking; previous studies in trees have only reported direct carryover effects or provenance-nursery environment interactions for a maximum of four years after transplantation/removal of the treatment (Dhar et al., 2014; Ewers et al., 1999). Both effects may have significant impacts on the survivorship of trees after planting, and hence on overall success of tree planting schemes. As many countries around the world are now engaged in major scaling up of tree planting to fulfil Net Zero carbon and biodiversity commitments, (e.g. the Bonn Challenge, Trillion Tree Campaign and the UK’s Nature for Climate Fund (Defra, UK Government, 2021)), an evaluation of the effect is overdue.

Scots pine is the most widely distributed pine species in the world with a native range extending from the Atlantic to the Pacific Oceans and from above the Arctic circle to the Mediterranean. It is an important foundation species in many Eurasian ecosystems, including the Caledonian pinewoods in Scotland, but it is also an economically valuable source of timber and is widely commercially planted. In the UK, plants for both commercial and conservation planting are grown from seed in domestic nurseries or, rarely, imported from nurseries in Europe (Richard Whittet, pers. comm., 2022). Seed is either sourced directly from native forests or from seed orchards which themselves were derived from native forests (Lee, 2002). Predominantly, plants are grown for around two years and supplied as bare root saplings (rather than containerised) before being transplanted (Whittet et al., 2016). There have been numerous studies that have identified significant differences in phenotypic traits among native UK Scots pine populations and families, even across the relatively small scale of the native range (Donnelly et al., 2018, 2016; Perry et al., 2016; Salmela et al., 2011). This suggests local adaptation to specific environmental conditions over a relatively small spatial scale, most likely caused by the significant spatial heterogeneity in climate across the native range. It is unknown whether the dynamics and persistence of carryover effects are affected by adaptive divergence among populations.

The impact of different nursery environments on growth and development and the strength and persistence of carryover effects may depend on the conditions in the nursery, the similarity of the environments at the nursery and planting site, and the similarity of the environments at the nursery and the seedling’s site of origin. This study aimed to evaluate carryover effects as long-term differences in the growth and phenology of trees due to their nursery environment. We used a long term study in which plants had been sourced from common origins, raised in different nurseries and planted in common garden trials (Beaton et al., 2022). We hypothesised that trees may show carryover effects either because they are differently provisioned in their early growth, because they acclimatise to environments during early growth, or because they are already genetically locally adapted to particular conditions. We propose three hypotheses to test these possible scenarios: the ‘provisioning hypothesis’; the ‘acclimation hypothesis’ and the ‘local adaptation hypothesis’

1. The *provisioning hypothesis* expects that trees from nurseries with less limiting environments (for example, a longer growing season or warmer temperatures during the growing season) will experience higher provisioning prior to transplantation and will have an advantage over trees from nurseries with more limiting environments, enabling them to grow more rapidly once transplanted (Gömöry et al., 2015)
2. The *acclimation hypothesis* expects that trees grown in nurseries local to the transplantation site will be acclimated to the environment at the point of transplantation and will therefore grow more rapidly than those which were not acclimated at the nursery stage, as acclimation will allow them to avoid or reduce transplantation shock (impaired growth or mortality after transplantation (Close et al., 2005).
3. The *local adaptation hypothesis* expects that trees grown in nurseries local to their site of origin and local to the transplantation site will be adapted and acclimated to the environment and will therefore grow more rapidly than those which were not adapted and/or not acclimated at the nursery stage.

## Method

### Source of plant material

A long term multisite common garden Scots pine trial was established in Scotland using seed collected from 21 populations from across the native range in Scotland (Figure 1) in March 2007 and is described in detail by Beaton et al., (2022). Briefly, seed was germinated and grown in an unheated glasshouse at the James Hutton Institute, Aberdeen (latitude 57.133214, longitude -2.158764) in June 2007. After around six weeks of growth, eight seedlings per family were transferred to each of three nursery sites (Figure 1): outdoors at Inverewe Gardens in the Scottish Highlands (NW, latitude 57.775714, longitude -5.597181); outdoors at the James Hutton Institute in Aberdeen (NE); in an unheated glasshouse at the James Hutton Institute in Aberdeen (NG). In 2012 the trees were transplanted to one of three field sites (Figure 1): Yair in the Scottish Borders (FS, latitude 55.603625, longitude -2.893025, planted in October 2012); Glensaugh (FE, planted in Spring 2012); and Inverewe (field site in the west of Scotland, planted in Spring 2012). Nursery and field site nomenclature follows the rule ‘XY’ where X refers to whether the site is a nursery (N) or field (F) site; and Y refers to the location of the site: G is the glasshouse, E is the east of Scotland, W is the west of Scotland and S is the south of Scotland. The two outdoor nurseries (NE and NW) were within the native range of Scots pine in Scotland and have distinct climates (Beaton et al., 2022). Two of the field sites were proximal to the outdoor nurseries (FE to NE; FW to NW), the third is outwith the current native range of Scots pine in Scotland (FS).

**Figure 1.**
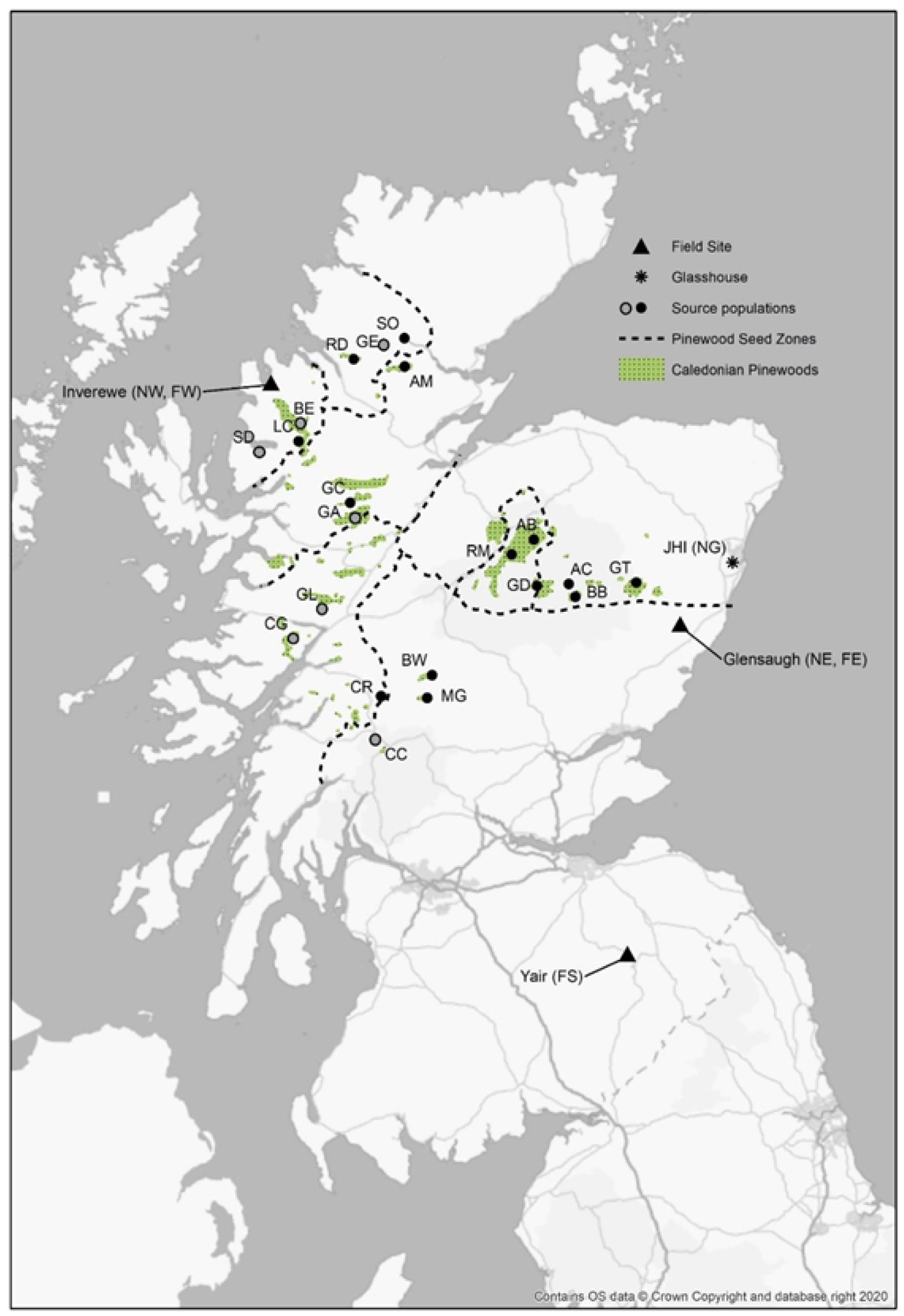
Location of the 21 source populations sampled for the multisite experimental trial; letter codes match those in Table S1. The subset of populations used in this study are in black, those not included are in grey. Also shown are James Hutton Institute (JHI), location of eastern nursery (NE) and glasshouse (NG); three field sites - Inverewe (FW, also location of western nursery: NW), Glensaugh (FE) and Yair (FS).

When the three field sites (FE, FW and FS) were planted in 2012, the experimental design was balanced and each site contained trees raised in its pre-determined single corresponding nursery (FE:NE, FW:NW and FS:NG). The only exception to this was a single tree at FW that was raised in NE. However, due to mortality at the field sites after transplantation, particularly at FW, two separate waves of ‘beating up’ were performed: in April 2013, four trees from NG were transplanted to FE and 81 trees from NE were transplanted to FW; then in October 2013, 132 trees from NG were transplanted to NW. This unforeseen modification in the experimental design provided an opportunity to assess the effect of the nursery environment on long term trait variation following transplantation.

In order to identify carryover effects, trait means relating to growth and phenology were compared for groups of trees that were planted at FW but which had been grown in NE, NG or NW. We carefully selected trees to form a dataset that was genetically balanced to allow comparisons, as follows. Across 14 of the populations (Figure 1; Table S1) planted at FW, groups were identified that had been grown at each nursery. As not all families were present within each group, it was not possible to have the same families present in all nursery groups, so three ‘family composition groups’ were formed (Table S2) such that common populations could be compared whilst keeping genetic variability within population consistent. Group A were grown in NE, Group B were grown in NG and Group C were grown in NW. The family composition groups were then used to select trees from the same families planted at either FE and FS for comparison. To check whether the difference in families among the Groups affected our results, trait means were compared for Groups A, B and C at FS and FE (Table S3a), where complete sampling was possible.

To test the *provisioning* and *acclimation hypotheses*, trait means relating to growth were compared to assess whether growth was higher for trees that were originally grown in a nursery with an environment that provided high levels of provisioning (i.e. NG, supporting the provisioning hypothesis) or in a nursery environment that allowed acclimation (i.e. NW, supporting the acclimation hypothesis) prior to transplantation.

To test the *local adaptation hypothesis*, trait means relating to growth were compared to assess whether growth was higher for trees that were raised and planted in areas to which they were local. For trees planted at NE and NW (Groups A and C, respectively), populations from the east (GT and BB) and west (LC and GC) of Scotland (Table S1) were selected and trees were assigned to one of three different groups depending on the locality of their source population, nursery and field site (Figure 1). Trees were classed as “Locally adapted and nursery acclimated” if their origin was local to both the nursery and the field site (i.e. NE-FE for populations from the east; NW-FW for populations from the west). Trees were classed as “Nursery acclimated” if their origin was not local to their nursery site but their planting site was local to their nursery (i.e. NE-FE for populations from the west; NW-FW for populations from the east). Trees were classed as “Nursery mismatched” if they were grown in nurseries which were nonlocal to the planting site (i.e. NE-FW for populations from both the east and west). The local adaptation hypothesis was supported if growth was higher for trees which were ‘Locally adapted and nursery acclimated’ compared to those from ‘Nursery acclimated’ or ‘Nursery mismatched’ groups.

### Phenotype assessments

Trait data were taken from 2014, the first full year all trees could be compared, until 2022. Measurement protocols are detailed in Beaton et al., (2022) and the associated datasets are publicly available (Perry et al., 2022). Briefly, absolute tree height was measured after the end of each growing season in each year. Annual height increment was estimated as the increase in height from one year to the next. Relative growth rate was expressed as a percentage, dividing annual increment by the absolute height in the previous year x 100. Absolute height and stem diameter were found to be highly significantly correlated in all years (Perry et al., 2024) but, as the ratio of these two traits has important implications for tree form, height to diameter ratio was also estimated for each year.

Phenology assessments were performed in spring at each field site from 2015 to 2022, apart from 2020 due to the Covid-19 pandemic. Each tree was assessed for budburst timing, considered as the time taken for needles to appear on the new shoot: assessments were performed at weekly intervals from early spring until budburst was complete. To allow comparisons within and among sites and years, variation in timing of budburst is considered relative to growing degree days (GDD). The growing degrees were estimated for each day from 1 January of each year as the number of degrees above 5 °C at each site, then GDD was estimated by summing the growing degrees for each period from 1 January to the date of budburst.

### Environmental variables

Budburst phenology for trees grown at each of the nurseries and planted at FW was compared to the mean monthly temperature for May (when trees were progressing through budburst) for 2015-2022. The closest weather station to FW was Poolewe (0.72 miles). Where Poolewe data were missing they were replaced with measurements from the next closest weather station: Aultbea No 2 (4.24 miles).

### Statistical analysis

One-way ANOVAs, with Tukey’s test for multiple pairwise comparisons, were performed in Minitab (version 21) to compare traits relating to growth (mean height, mean height increment, mean relative growth rate, mean height to diameter ratio) and phenology (mean GDD to budburst) to test hypotheses outlined in Figure 2.

**Figure 2.**
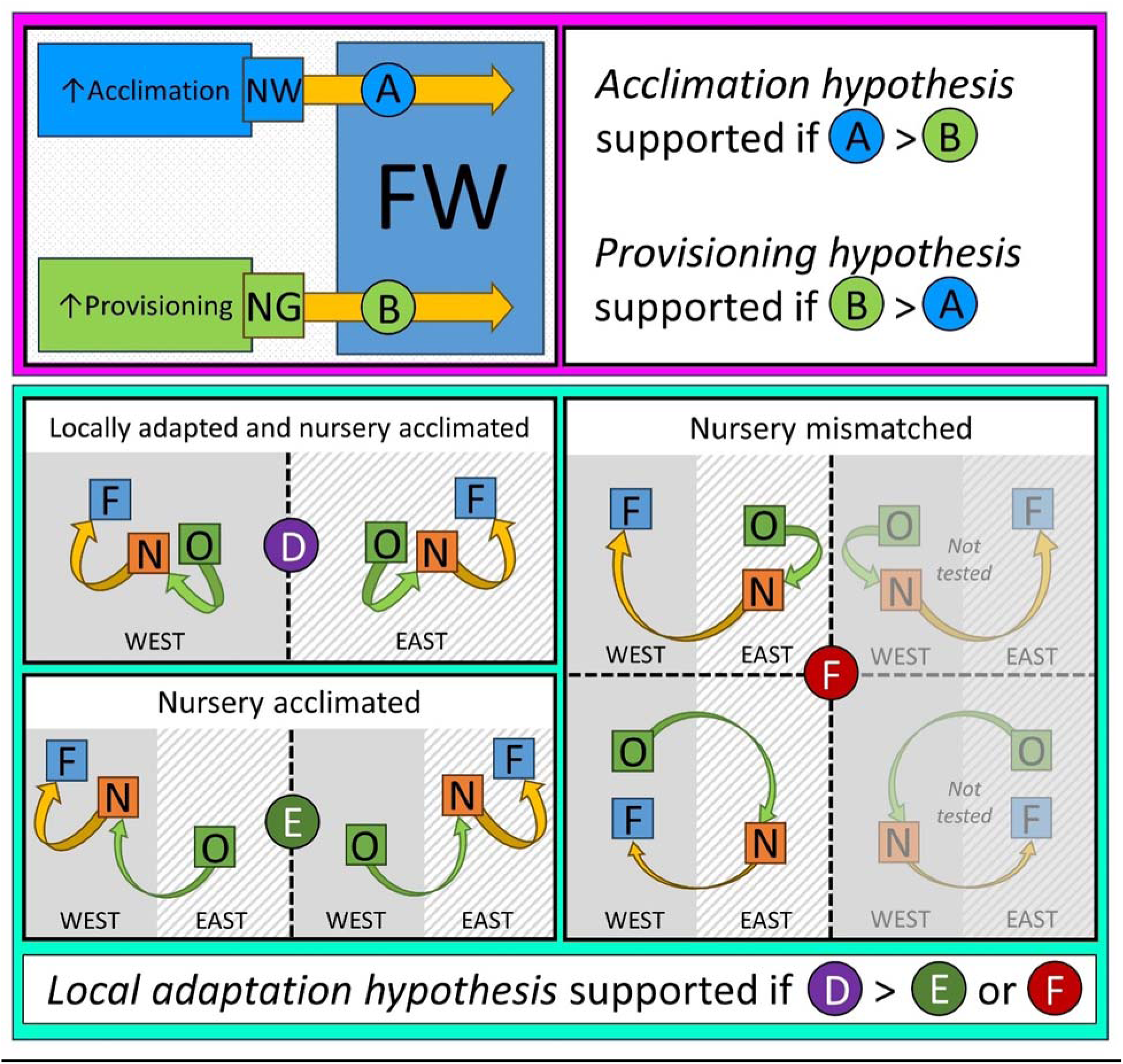
Schematic for testing hypotheses relating to carryover effects (acclimation hypothesis, provisioning hypothesis and local adaptation hypothesis). The acclimation and provisioning hypotheses (pink box) are tested by comparing trait variation for trees planted at the same site (FW) but grown in two different nursery scenarios: a local nursery which enabled acclimation prior to transplantation (scenario A: NW); or a high provisioning nursery (scenario B: NG). The local adaptation hypothesis (turquoise box) is tested by comparing trait variation for trees originating, grown and planted in a further three scenarios: trees planted in field sites local to the nursery site and local to their origin (scenario D: locally adapted and nursery acclimated); trees planted in field sites local to the nursery site but nonlocal to their origin (scenario E: nursery acclimated); and trees planted in field sites nonlocal to their nursery site (scenario F: nursery mismatched). Two other combinations are theoretically possible for scenario F, but were not tested due to a lack of trees – greyed out.

## Results

To check whether any differences observed might be due to among-family composition within the sampling groups, we evaluated groups of trees composed of the same families but that had been planted at the other field sites, FE and FS – i.e. where groups differed only in their family composition and not nursery environment (Table S3a).

These groups showed no significant differences in their means for growth (absolute or increment height and relative growth rate) or for GDD to budburst for any of the years assessed except: FS, height increment in 2016; and FE, GDD to budburst in 2022. We therefore assume that any differences detected among groups at FW can reliably be attributed to differences in the nursery environments they experienced.

### Extent and persistence of the carryover effect

Trees transplanted to FW that originated from different nurseries showed significant differences in growth and budburst phenology for many years after transplantation (Table S3b) indicating that carryover effects were strong and persistent.

In phenology, trees grown in NE burst bud after significantly fewer GDD than trees grown in NG and NW in three out of five years (Figure 3, Table S3b; although in 2018 only trees from NG and NE were significantly different for this trait). In years where mean GDD to budburst was significantly different among nursery groups (Figure 3), the average mean May temperature was much warmer (2016: 11.6 °C; 2017: 12.6 °C; 2018: 12.1 °C) than when it was not significantly different (2015: 8.8 °C; 2019: 9.4 °C, 2021: 9.1 °C). The exception to this was 2022 when the average mean May temperature was 11.1 °C but GDD to budburst was not significantly different among nursery groups, possibly indicating a waning of the carryover effect.

**Figure 3.**
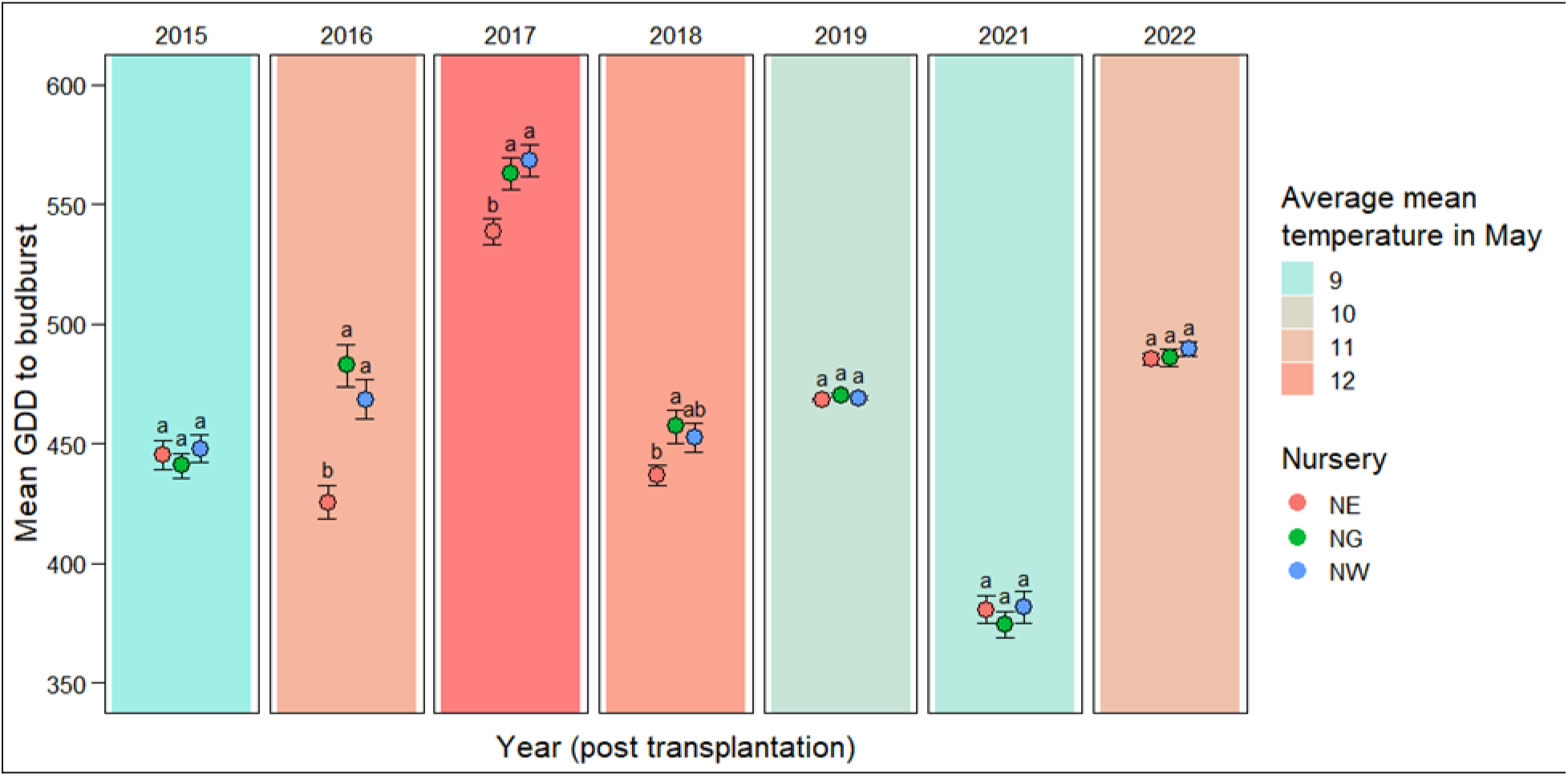
Mean growing degree days (GDD) to budburst for trees grown at each of three nurseries (NE, NG, NW) following transplantation to FW. Error bars indicate one standard error either side of the mean. Tukey’s post-hoc test for each year grouped nurseries with significantly different mean values (a, b): letters are shared among nurseries where mean values are not significantly different. Average mean temperature for May is indicated by colour of background shading each year: blue indicates colder temperatures and red indicates warmer temperatures.

### Testing the provisioning and acclimation hypotheses

To understand the dynamics underlying the carryover effects and to test the provisioning and acclimation hypotheses, the growth of trees from the different nurseries was compared to one another. Trees from NG, which we assumed was a high provisioning environment, were taller in the earliest years of field growth compared to trees grown in NW (in 2014: mean height NG group = 1218 mm; NW group = 874 mm). However, trees in the NG group subsequently showed poorer relative growth rates in all years compared to trees from NW. These findings support the acclimation hypothesis over the provisioning hypothesis. Annual height increments were significantly higher for trees from NW compared to those from nonlocal nurseries from 2014 to 2019 (Figure 4, Table S3b); after 2019 the mean values are higher (except for 2021) but not significantly.

**Figure 4.**
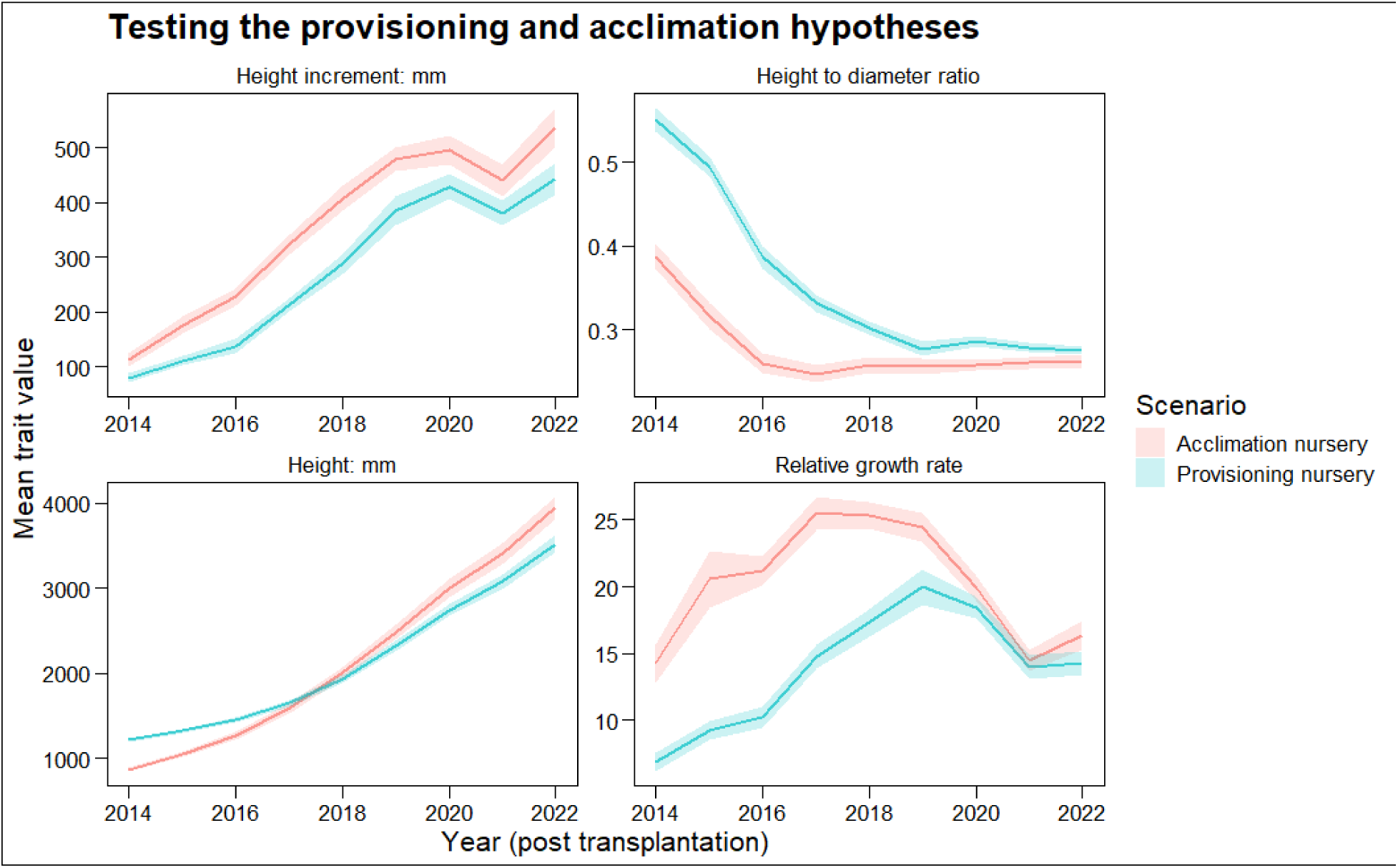
Variation in growth traits for trees grown at NG (provisioning nursery) and NW (acclimation nursery). Lines indicate trait means for each nursery from 2014 to 2022; shaded areas around lines indicate one standard error either side of the mean.

In 2014, the mean height to diameter ratio was much higher for trees from NG (0.55) than for trees from NW (0.39). Intriguingly, for trees from NW the ratio appeared to level out in 2016 at around 0.25 (Figure 4), but trees from NG took until 2019 to level out at around 0.28.

### Testing the local adaptation vs acclimation hypotheses

The effect of the locality of nurseries and field sites on growth traits was compared for populations from the east and west of Scotland. For both groups, mismatching the nursery and field sites led to significantly smaller trees (Figure 5; Table S3c) compared to trees that were grown and planted in nursery and field sites local to their site of origin (“locally adapted and nursery acclimated”). This difference was much more pronounced for trees from the east of Scotland, for which mismatching produced consistently and significantly lower annual increment gains for up to five years after transplantation (Figure 5; Table S3c). The height to diameter ratio was also considerably larger for mismatched eastern trees compared to those that were either ‘nursery acclimated’ or ‘locally adapted and nursery acclimated’, until 2018 when the ratio between the traits levelled out. As was seen above in the difference in carryover effects, although the differences in growth rate reduced or disappeared within a few years of transplantation, the effect on the absolute height, as a cumulative trait, was still apparent even 10 years after transplantation. In the western group ‘locally adapted and nursery acclimated’ trees grew tallest, whilst in the eastern group ‘nursery acclimated’ trees grew tallest (Figure 5).

**Figure 5.**
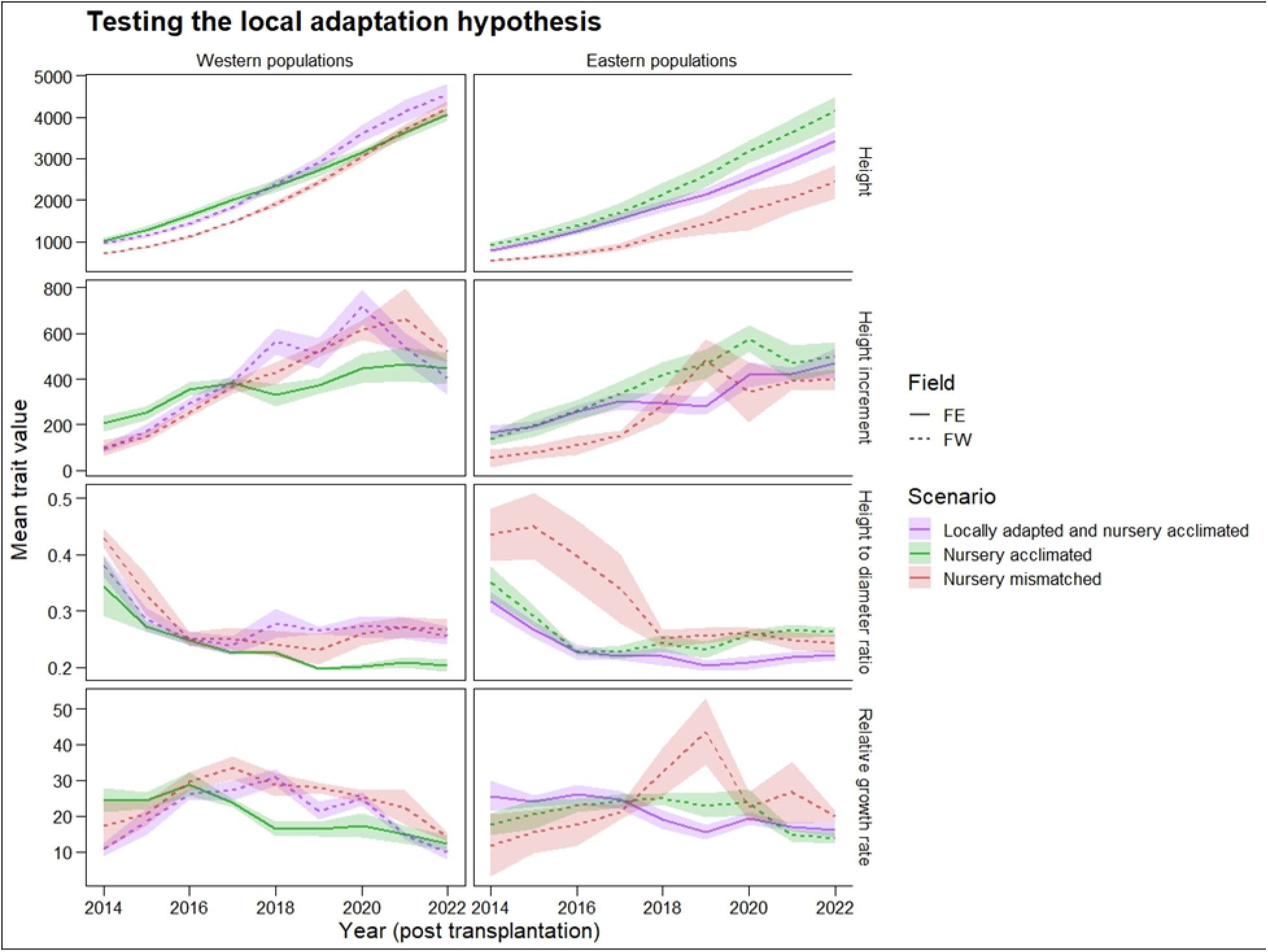
Testing the local adaptation hypothesis (see Figure 2) by comparing variation in mean growth traits from 2014 to 2022 for trees from the western (LC and GC) and eastern (BB and GT) groups. Trees were either: planted in field sites local to the nursery site and local to their origin (scenario D in Figure 2: ‘locally adapted and nursery acclimated’); planted in field sites local to the nursery site but nonlocal to their origin (scenario E, figure 2: ‘nursery acclimated’); planted in field sites nonlocal to their nursery site (scenario F, Figure 2: ‘nursery mismatched’). Shaded area around lines indicate one standard error either side of the mean. Field site is also indicated by line type (solid line: FE; dashed line: FW) to show growth differences among trees in different environments after transplantation.

The results from our study supported our *acclimation hypothesis*, with support for our *local adaptation hypothesis* in some environments. There was no support for the *provisioning hypothesis*, although future work with an increased number of test environments (both nursery and field) and with reciprocity built into the design would enable firmer conclusions to be drawn. Trees that were acclimated to the environment in the west of Scotland showed consistently higher values for growth traits in this environment, compared to trees which had not been acclimated but had received high levels of provisioning in their nursery. This effect was still apparent 10 years after transplantation and, although the difference in growth rates appears to have reduced, the cumulative effect of many years of growing at different rates means that the overall height of the trees is now markedly different. Trees grown in the east of Scotland showed similar patterns of acclimation: those planted locally to their nursery environment grew taller and had larger growth increments for almost every year recorded after transplantation, compared to those that were raised in the east but planted in the west of Scotland. In contrast, and despite their initial growth advantage, trees from the glasshouse nursery showed consistently lower growth compared to the acclimated trees. Close et al., (2005) suggest that trees that are not acclimated to their planting site environment remain stressed until acclimation occurs. This would imply that carryover effects are driven by a stress-induced disadvantage for those which are not acclimated to conditions (and conversely that acclimated trees gain a growth advantage due to a comparatively low stress response). Stress-induced epigenetic effects are becoming a well-recognised part of plant phenotypic variation (Mirouze and Paszkowski, 2011) and it seems possible that such an effect is operating here. Although we did not explicitly evaluate the mechanisms underlying the carryover effects, future analysis of the role of epigenetic modification in shaping them would be beneficial.

The allocation of resources to height or stem diameter, which was different among trees from different nurseries and varied over the duration of the study, highlighted the extent to which trees were acclimated to the outdoor environment and the impact that acclimation had on the subsequent growth of the trees. A proportionally greater investment in height may indicate an environment where increasing photosynthetic capacity provides a competitive advantage (King, 1990), whilst a greater investment in stem diameter is thought to indicate environments where more mechanical support and/or water-absorbing capacity provides a competitive advantage (Close et al., 2005; Munishi and Chamshama, 1994; Niklas, 1993). Exposure to wind, rain and snow, which trees experienced to differing degrees in outdoor nurseries and not at all in the glasshouse, can result in an increase in stem diameter as a proportion of height in order to increase the mechanical strength of the main stem (Claussen and Maycock, 1995; Close et al., 2005). Acclimation to these abiotic stresses had a demonstrable impact on height to stem diameter ratios. This trend was observable not only for trees that had been grown in the glasshouse, but also for trees grown in an outdoor nursery that did not match their planting site.

Significantly, carryover effects were also observed in budburst timing up to six years after transplantation to a common environment. Trees that had acclimatised to the cooler environment in the nursery in the east of Scotland required fewer GDD to reach budburst, especially when Spring temperatures were particularly warm. This was already apparent when trees were assessed in the nursery (Perry et al., 2024) but it is striking that the acclimatisation persisted into the field for several years. This implies that acclimatisation is enacted at a very early stage, undergoing little subsequent adjustment even if conditions change. Budburst initiation in cooler environments is triggered at a lower temperature sum threshold in comparison to warmer environments (Gömöry et al., 2015; Rötzer et al., 2004), so after transplantation to the common environment at FW, trees from NE burst bud earlier than those from NG and NW. Gömöry et al., (2015) reported similarly delayed budburst in *Picea abies* and *Larix decidua* whose early environment was in a warm nursery compared to those grown in a cold nursery, regardless of where they were later transplanted to. Another study reported significant carryover effects on phenology following transplantation for *Betula papyifera*, but only for one year after planting and the effect disappeared altogether after three years (Dhar et al., 2015). Trees which experienced early growth in cooler environments seem to show evidence of acclimation: they require a lower temperature sum for the initiation of budburst, which persists for many years after transplantation into a common environment. This has significant implications, although phenological mismatch with the environment can be negative (i.e. new shoots may be vulnerable to frost damage) or positive (i.e. an effectively longer growing season). It may also have knock-on effects on reproductive receptivity (Whittet et al., 2017). Although in our study the effect seems to have disappeared in the most recent years, the apparent association of the timing differences with particularly warm May temperatures means that effects may still re-appear in the future if warmer conditions occur.

Local adaptation also had a noticeable effect on growth, with trees from the west of Scotland showing higher plasticity in their response to varying nursery and field environments than trees from the east of Scotland. They were also less impacted by mismatching of the nursery and field sites. It seems that although nursery acclimation was more important than local adaptation when considering the magnitude of impact on trait variation, the effect was evident only in some environments. Alternatively, the difference in plasticity itself may constitute adaptive divergence (Chevin and Lande, 2011; Linhart and Grant, 1996). It is often reported that planting locally-sourced trees has benefits relating to their expected adaptation to the environment (Jones et al., 2001; O’Brien and Krauss, 2008; Sackville Hamilton, 2001), and local adaptation is the basis for much seed zoning policy (Breed et al., 2013). However, the effects and impacts of local adaptation are not usually considered in the context of an interaction between the nursery and planting environments and this effect would benefit from further research.

## Conclusion

This study highlights the importance of early environment in the subsequent growth and development of trees following transplantation. Given the increasing scale of tree planting around the world, as part of strategies to achieve net zero carbon emissions and to address the biodiversity crisis, attention needs to be paid to factors that influence the success or failure of tree in the field. Given their observed extent and persistence, it is clear we need much greater understanding of the potential costs and benefits of carryover effects arising from the nursery environment across a wide range of species. To better understand the interactions between nurseries, field sites and sites of origin, a carefully designed experiment would be necessary. Ideally, this should be a fully reciprocal design across nursery and field sites, with scope to be monitored for multiple years and potentially long-term. It would also be interesting to test how long trees need to be grown in an environment for acclimation to occur: in this study, trees were grown in their early environment for nearly six years, which may have resulted in a more pronounced effect than if they had been transplanted after only one or two years (as is commonplace in commercial nurseries).

## Acknowledgements

We thank Stewart Mackie and staff at Forest and Land Scotland (Selkirk Office), the Forestry Commission, National Trust for Scotland, and Donald Barrie (Glensaugh) for their help in locating, establishing and securing access to sites.

## Data availability statement

The data underlying this article are available from the Environmental Information Data Centre (EIDC, https://eidc.ac.uk), at https://doi.org/10.5285/1c9367fb-ea87-47a1-8257-d9fed54215e7

