## Supplementary materials for "Significant and persistent carryover effects in Scots pine"

Table S1. Locations and basic environmental data for the subset of populations from the multisite trial which were included in this study

| Population | Lat. (N) | Long. (W) | Alt. (m) | GSL | GDD | FMT (°C) | JMT (°C) |
| --- | --- | --- | --- | --- | --- | --- | --- |
| Abernethy (AB) | 57.21 | 3.61 | 311-370 | 211 | 990 | 1.1 | 12.7 |
| Allt Cul (AC) | 57.04 | 3.35 | 435-512 | 145 | 513 | -1.0 | 10.4 |
| Amat (AM) | 57.87 | 4.60 | 39-201 | 214 | 892 | 1.2 | 12.3 |
| Ballochbuie (BB) | 56.98 | 3.30 | 421-531 | 116 | 446 | -1.7 | 9.5 |
| Black Wood of Rannoch (BW) | 56.68 | 4.37 | 250-321 | 254 | 1138 | 2.1 | 13.5 |
| Crannach (CR) | 56.58 | 4.68 | 258-338 | 231 | 1019 | 1.8 | 12.6 |
| Glen Cannich (GC) | 57.35 | 4.95 | 182-381 | 212 | 778 | 1.0 | 11.7 |
| Glen Derry (GD) | 57.03 | 3.58 | 426-493 | 168 | 593 | -0.5 | 11.3 |
| Glen Tanar (GT) | 57.02 | 2.86 | 289-422 | 235 | 1105 | 2.2 | 13.6 |
| Loch Clair (LC) | 57.56 | 5.36 | 98-166 | 277 | 1253 | 3.4 | 13.7 |
| Meggernie (MG) | 56.58 | 4.35 | 254-385 | 223 | 916 | 1.1 | 12.0 |
| Rhidorroch (RD) | 57.89 | 4.98 | 138-220 | 221 | 840 | 1.5 | 11.6 |
| Rothiemurchus (RM) | 57.15 | 3.77 | 295-329 | 224 | 1087 | 1.4 | 13.1 |
| Strath Oykel (SO) | 57.98 | 4.61 | 35-160 | 257 | 1276 | 2.7 | 14.0 |

Alt: altitudinal range sampled within each population; Average (1961-2000) climate variables from UK Met Office (Perry and Hollis 2005): GSL, growing season length (period bounded by daily mean temperature > 5 °C for > 5 consecutive days and daily mean temperature < 5 °C for > 5 consecutive days (after 1 July); GDD, growing degree days (the mean number of degrees by which the air temperature has gone above 5 °C calculated day by day and summed over the year); FMT and JMT, February and July mean temperatures (°C).

Table S2. Counts of families representing each family composition group (A, B, C). Family codes consist of population code (first two letters: see Table S1) and number.

| Family | A | B | C |  | Family | A | B | C |  | Family | A | B | C |  | Family | A | B | C |
| --- | --- | --- | --- | --- | --- | --- | --- | --- | --- | --- | --- | --- | --- | --- | --- | --- | --- | --- |
| AB10 |  | 1 |  |  | BB2 |  |  | 1 |  | GC9 |  |  | 2 |  | MG8 |  |  | 1 |
| AB3 | 1 |  |  |  | BB5 |  | 1 | 1 |  | GD1 |  |  | 1 |  | MG9 |  | 1 | 1 |
| AB4 | 1 |  | 1 |  | BB7 |  | 1 |  |  | GD10 | 2 | 1 |  |  | RD1 | 1 | 1 |  |
| AB5 | 1 |  | 1 |  | BB8 |  | 1 |  |  | GD2 |  | 1 |  |  | RD10 |  | 1 |  |
| AB6 |  | 1 |  |  | BB9 | 2 |  |  |  | GD5 |  | 1 | 1 |  | RD2 | 1 |  |  |
| AB9 |  | 1 | 1 |  | BW10 |  | 1 | 1 |  | GD6 |  |  | 1 |  | RD3 | 1 |  | 1 |
| AC1 |  |  | 1 |  | BW2 |  | 1 | 2 |  | GD7 | 1 |  |  |  | RD5 |  |  | 1 |
| AC2 |  | 1 |  |  | BW3 | 1 |  |  |  | GT1 |  | 1 |  |  | RD7 |  | 1 |  |
| AC3 | 1 |  | 1 |  | BW4 | 1 |  |  |  | GT2 | 1 |  |  |  | RD8 |  |  | 1 |
| AC5 | 1 | 1 |  |  | BW6 | 1 | 1 |  |  | GT5 |  |  | 1 |  | RM1 | 1 |  |  |
| AC6 |  | 1 |  |  | CR2 | 1 | 1 |  |  | GT6 |  | 1 | 1 |  | RM10 |  | 1 |  |
| AC7 | 1 |  |  |  | CR3 |  |  | 1 |  | GT7 | 1 | 1 | 1 |  | RM3 | 1 | 1 | 1 |
| AC9 |  |  | 1 |  | CR4 | 1 |  |  |  | GT9 | 1 |  |  |  | RM5 | 1 | 1 |  |
| AM1 |  |  | 1 |  | CR5 | 1* | |  |  | LC1 |  |  | 1 |  | RM7 |  |  | 2 |
| AM10 |  | 1 |  |  | CR6 |  | 1 |  |  | LC2 |  | 1 |  |  | SO10 |  | 1 | 1 |
| AM3 |  |  | 1 |  | CR7 |  | 1 | 1 |  | LC4 | 1 | 1 |  |  | SO2 |  | 1 |  |
| AM4 |  | 1 |  |  | CR9 |  |  | 1 |  | LC5 |  | 1 | 1 |  | SO3 | 1 |  | 1 |
| AM5 | 1 |  |  |  | GC1 |  | 1 | 1 |  | LC7 | 2 |  | 1 |  | SO4 | 1 |  |  |
| AM6 | 1 |  |  |  | GC10 |  | 1 |  |  | MG10 |  | 1 |  |  | SO5 |  | 1 |  |
| AM8 |  | 1 | 1 |  | GC2 | 1 |  |  |  | MG2 | 1 | 1 |  |  | SO6 |  |  | 1 |
| AM9 | 1 |  |  |  | GC6 | 1 | 1 |  |  | MG3 | 1 |  |  |  | SO7 | 1 |  |  |
| BB1 | 1 |  | 1 |  | GC8 | 1 |  |  |  | MG5 | 1 |  | 1 |  |  |  |  |  |

Table S3a-c. Comparing mean trait values (HA: absolute height; HI: height increment; HR: relative growth rate; growing degree days to budburst: BB) for each year assessed (no assessment: ‘-’) for trees grown at different nurseries (NE; NW; NG) and transplanted to different field sites (FE; FW; FS) using family composition groups (‘Family’) A-C (see Table S2). Tukey’s tests for multiple pairwise comparisons are reported: groups that do not share a letter are significantly different. Different combinations of nursery and field site are included depending on the comparisons made. Years assessed: 2014 (‘14’) to 2022 (‘22’).

Table S3a. Checking for differences among traits caused by family composition of nursery groups (see Table S1).

|  |  |  |  |  | Year | | | | | | | | |
| --- | --- | --- | --- | --- | --- | --- | --- | --- | --- | --- | --- | --- | --- |
| Trait | Nursery | Field | Family | N | 14 | 15 | 16 | 17 | 18 | 19 | 20 | 21 | 22 |
| *Field site: FS* | | | | | | | | | | | | | |
| HA | NG | FS | A | 41 | a | a | a | a | a | a | a | a | a |
|  | NG | FS | B | 42 | a | a | a | a | a | a | a | a | a |
|  | NG | FS | C | 42 | a | a | a | a | a | a | a | a | a |
| HI | NG | FS | A | 41 | - | a | a | a | a | a | a | a | a |
|  | NG | FS | B | 42 | - | a | b | a | a | a | a | a | a |
|  | NG | FS | C | 42 | - | a | ab | a | a | a | a | a | a |
| HR | NG | FS | A | 41 | - | a | a | a | a | a | a | a | a |
|  | NG | FS | B | 42 | - | a | a | a | a | a | a | a | a |
|  | NG | FS | C | 42 | - | a | a | a | a | a | a | a | a |
| BB | NG | FS | A | 41 | - | a | a | a | a | a | - | a | a |
|  | NG | FS | B | 42 | - | a | a | a | a | a | - | a | a |
|  | NG | FS | C | 42 | - | a | a | a | a | a | - | a | a |
| *Field site: FE* | | | | | | | | | | | | | |
| HA | NE | FE | A | 42 | a | a | a | a | a | a | a | a | a |
|  | NE | FE | B | 42 | a | a | a | a | a | a | a | a | a |
|  | NE | FE | C | 42 | a | a | a | a | a | a | a | a | a |
| HI | NE | FE | A | 42 | a | a | a | a | a | a | a | a | a |
|  | NE | FE | B | 42 | a | a | a | a | a | a | a | a | a |
|  | NE | FE | C | 42 | a | a | a | a | a | a | a | a | a |
| HR | NE | FE | A | 42 | a | a | a | a | a | a | a | a | a |
|  | NE | FE | B | 42 | a | a | a | a | a | a | a | a | a |
|  | NE | FE | C | 42 | a | a | a | a | a | a | a | a | a |
| BB | NE | FE | A | 42 | - | a | a | a | a | a | - | a | a |
|  | NE | FE | B | 42 | - | a | a | a | a | a | - | a | b |
|  | NE | FE | C | 42 | - | a | a | a | a | a | - | a | ab |

Table S3b. Comparing trees grown in different nurseries (NE, NG, NW) and planted at the same field site (FW): used to test the acclimation/provisioning hypotheses (Figure 2)

|  |  |  |  |  | Year | | | | | | | | |
| --- | --- | --- | --- | --- | --- | --- | --- | --- | --- | --- | --- | --- | --- |
| Trait | Nursery | Field | Family | N | 14 | 15 | 16 | 17 | 18 | 19 | 20 | 21 | 22 |
| HA | NE | FW | A | 42 | c | c | c | b | b | b | b | b | c |
|  | NG | FW | B | 42 | a | a | a | a | a | a | a | a | b |
|  | NW | FW | C | 42 | b | b | b | a | a | a | a | a | a |
| HI | NE | FW | A | 42 | b | b | b | b | b | ab | a | a | a |
|  | NG | FW | B | 42 | b | b | b | b | b | b | a | a | a |
|  | NW | FW | C | 42 | a | a | a | a | a | a | a | a | a |
| HR | NE | FW | A | 42 | b | a | b | a | a | a | a | a | a |
|  | NG | FW | B | 42 | b | b | c | b | b | b | b | b | a |
|  | NW | FW | C | 42 | a | a | a | a | a | b | ab | b | a |
| BB | NE | FW | A | 42 | - | a | b | b | b | a | - | a | a |
|  | NG | FW | B | 42 | - | a | a | a | a | a | - | a | a |
|  | NW | FW | C | 42 | - | a | a | a | ab | a | - | a | a |

Table S3c. Comparing locality of trees to their nursery and/or field sites: used to test the local adaptation hypothesis (Figure 2). Eastern populations: NE+FE, local and nursery acclimated; NW+FW, nursery acclimated; NE+FW, nursery mismatched. Western populations: NW+FW, local and nursery acclimated; NE+FE, nursery acclimated; NE+FW, nursery mismatched.

|  |  |  |  |  | Year | | | | | | | | |
| --- | --- | --- | --- | --- | --- | --- | --- | --- | --- | --- | --- | --- | --- |
| Trait | Nursery | Field | Family | N | 14 | 15 | 16 | 17 | 18 | 19 | 20 | 21 | 22 |
| *Eastern populations: BB and GT* | | | | | | | | | | | | | |
| HA | NW | FW | C | 6 | a | a | a | a | a | a | a | a | a |
|  | NE | FE | C | 6 | a | a | a | a | ab | ab | ab | ab | ab |
|  | NE | FW | A | 6 | b | b | b | b | b | b | b | b | b |
| HI | NW | FW | C | 6 | a | a | a | a | a | a | a | a | a |
|  | NE | FE | C | 6 | a | a | ab | ab | a | a | a | a | a |
|  | NE | FW | A | 6 | a | a | b | b | a | a | a | a | a |
| HR | NW | FW | C | 6 | a | a | a | a | a | b | a | a | a |
|  | NE | FE | C | 6 | a | a | a | a | a | b | a | a | a |
|  | NE | FW | A | 6 | a | a | a | a | a | a | a | a | a |
| BB | NW | FW | C | 6 | - | a | a | a | a | a | - | b | b |
|  | NE | FE | C | 6 | - | b | a | a | b | b | - | a | a |
|  | NE | FW | A | 6 | - | a | a | a | a | a | - | b | b |
| Western populations: LC and GC | | | | | | | | | | | | | |
| HA | NW | FW | C | 6 | a | a | a | a | a | a | a | a | a |
|  | NE | FE | C | 6 | a | a | a | a | a | a | ab | a | a |
|  | NE | FW | A | 6 | b | b | b | b | b | a | b | a | a |
| HI | NW | FW | C | 6 | b | a | ab | a | a | a | a | a | a |
|  | NE | FE | C | 6 | a | a | a | a | b | a | b | a | a |
|  | NE | FW | A | 6 | ab | a | b | a | ab | a | ab | a | a |
| HR | NW | FW | C | 6 | b | a | a | ab | a | ab | a | a | a |
|  | NE | FE | C | 6 | a | a | a | b | b | b | a | a | a |
|  | NE | FW | A | 6 | ab | a | a | a | a | a | a | a | a |
| BB | NW | FW | C | 6 | - | a | a | a | a | a | - | a | a |
|  | NE | FE | C | 6 | - | b | a | a | a | b | - | b | b |
|  | NE | FW | A | 6 | - | a | a | a | a | a | - | a | a |
